# Transfer of symbolic numeral adaptation across eyes and hemifields

**DOI:** 10.64898/2026.03.10.710478

**Authors:** Ayuka Nakamura, Junxiang Luo, Isao Yokoi, Hiromasa Takemura

## Abstract

Visual perception of symbolic numerals is essential for everyday tasks; however, the neural and perceptual mechanisms underlying this ability remain unclear. Partially occluded digital numerals can elicit bistable perception, and adaptation to symbolic numerals alters the perception of these ambiguous stimuli. We aimed to examine how symbolic numeral adaptation is related to hierarchical visual processing by testing its interocular and interhemifield transfer. Experiment 1 tested interocular transfer by presenting the test stimulus to either the same or opposite eye as the adaptation stimulus. Experiment 2 assessed interhemifield transfer by presenting the test stimulus to either the same or opposite hemifield as the adaptation stimulus. Experiment 3 examined the interhemifield transfer of adaptation confined to the upper parts of digital numerals. Our results showed that adaptation to digital numerals induced shifted perceptual interpretations that transferred across eyes. In addition, we found that adaptation to digital numerals induced a relatively small but statistically significant interhemifield transfer. In contrast, adaptation restricted to the upper parts of digital numerals showed no significant interhemifield transfer. These findings suggest that the perceptual interpretation of symbolic numerals involves visual processing stages that integrate information across the eyes and hemifields.

## Introduction

Understanding the visual mechanisms underlying numerical perception is a fundamental goal in vision science. Psychophysical and physiological studies have investigated this question using nonsymbolic numerical stimuli, such as dot arrays. These studies have demonstrated the numerosity adaptation effect (Aagten-Murphy & Burr, 2016; Anobile et al., 2018; Burr & Ross, 2008), identified neurons selectively tuned to nonsymbolic numerical information (Nieder & Dehaene, 2009), and revealed cortical maps selective for nonsymbolic numeral information (Harvey et al., 2013).

However, another essential aspect of human numerical cognition is the perceptual recognition of symbolic numerals, such as Arabic digits, which is an indispensable skill for everyday tasks, including reading clocks and interpreting prices. Previous studies have investigated how symbolic numbers are represented in the occipitotemporal and inferotemporal cortices to elucidate the neural mechanisms underlying symbolic numeral processing (Cai et al., 2023; Conrad et al., 2023; Grotheer et al., 2016; Shum et al., 2013). However, the stage within the visual hierarchy at which symbolic numeral processing occurs remains unclear.

In the visual system, the hierarchical stages of processing are closely associated with how information from each eye and visual hemifield is represented and integrated in the cortex. In humans and nonhuman primates, retinal inputs reach the primary visual cortex (V1) through early white matter pathways (Takemura et al., 2024). At this early stage, signals from the two eyes remain segregated in ocular dominance columns (Cheng et al., 2001; Horton et al., 1990), and responses are largely restricted to stimuli presented in the contralateral visual hemifield, reflecting the organization of early retinogeniculate pathways (Amano et al., 2009; Tootell et al., 1998). At later stages, these monocular signals are progressively integrated to support binocular processing (Goncalves et al., 2015; Preston et al., 2008; Tsao et al., 2003), and visual field representations of higher-level visual areas expand to include information from both visual hemifields (Mackey et al., 2017; Tootell et al., 1998), most likely because of interhemispheric information exchange via the corpus callosum (Dougherty et al., 2005; Van Essen & Zeki, 1978). Therefore, assessing whether a given perceptual phenomenon depends on eye- or hemifield-specific representations provides a principled approach for inferring the hierarchical stage at which the underlying visual processing emerges.

A recent psychophysical study (Luo et al., 2024) reported a phenomenon that may provide insight into how symbolic numeral information is processed within a visual hierarchy. The authors demonstrated that a partially occluded digital numeral could elicit bistable perception and that adaptation to one numeral induced a perceptual bias toward the unadapted interpretation. A similar though weaker adaptation effect was observed when participants adapted to the visual elements corresponding to parts of the numerals. These findings provide psychophysical evidence of the mechanisms that process visual number forms and complex shapes, which may correspond to different stages of the visual hierarchy. However, the specific locus within the visual hierarchy underlying this phenomenon remains unclear.

Thus, this study aimed to identify the stage of visual information processing underlying the symbolic numeral adaptation effect reported by Luo et al. (2024). To this end, we adopted the approach commonly used in psychophysical studies that quantifies the transfer of adaptation effects when the test and adaptation stimuli are presented to different eyes or visual hemifields (Mitchell & Ware, 1974; Movshon et al., 1972; Nakayama et al., 2024; Nishida et al., 1994; Palmer & Clifford, 2022). Because eye- and hemifield-specific information is progressively integrated along the visual hierarchy, the degree of interocular and interhemifield transfer can provide a principal means of inferring the stage of visual processing crucial for this perceptual bias.

In Experiment 1, we manipulated whether adaptation (normal digital numerals) and test stimuli (occluded digital numerals) were presented to the same or opposite eye to assess the relative contributions of monocular and binocular processing to the symbolic numeral adaptation effect. In Experiment 2, we varied whether adaptation occurred in the same or opposite visual hemifield as the test stimulus, allowing us to examine whether the effect occurs before or after the interhemispheric integration of visual information. In Experiment 3, we tested interhemifield transfer when adaptation stimuli consisted only of the upper parts of digital numerals to evaluate whether adaptation at a feature-based (shape component) processing stage engages hierarchical stages similar to whole number form adaptation. Across these experiments, we quantified the extent to which the symbolic numeral adaptation effect is transferred across the eyes and hemifields, thereby providing converging evidence regarding its locus within the visual hierarchy.

## Methods

### Participants

Twenty-six adults participated in Experiment 1 (20 females and 6 males; age range, 20–50 years; mean age, 38.15 years), 24 adults participated in Experiment 2 (16 females and 8 males; age range, 18–48 years; mean age, 32.36 years), and 23 adults participated in Experiment 3 (13 females and 10 males; age range, 20–50 years; mean age, 33.17 years). All the participants had normal or corrected-to-normal visual acuity. All the participants were recruited from the Okazaki area through the participant pool of the National Institute for Physiological Sciences.

This study was approved by the Ethics Committee of the National Institutes of Natural Sciences (protocol number: EC01-64). The participants provided written informed consent after receiving information about their right to withdraw at any time, the voluntary nature of participation, experimental procedures, and permission to share anonymized data. Before the main experiment, the participants completed a screening battery to assess their visual acuity, astigmatism, and binocular stereopsis.

One participant in Experiment 2 withdrew before completing all trials; therefore, analyses on Experiment 2 were performed using data from the remaining 23 participants (16 females and 7 males; age range, 18–48; mean age, 33.09 years).

### Apparatus

Visual stimuli were presented on a 32-inch Display++ LCD monitor (Cambridge Research Systems Ltd., Rochester, UK) with a spatial resolution of 1,920 × 1,080 pixels and a refresh rate of 120 Hz. The luminance output of the display was gamma-linearized. The visual stimuli were generated on a Dell Precision 5820 workstation (Round Rock, TX, USA) running Ubuntu 20.04 LTS.

The participants’ heads were stabilized using a custom-built chin rest positioned 65 cm from the monitor to maintain a constant viewing distance. Fixation was monitored using an eye-tracking system (LiveTrack Lightning; Cambridge Research Systems Ltd., Rochester, UK), with the camera positioned 18 cm in front of the eyes. The system continuously tracked the positions of both eyes in real time. The participants reported their perceptual responses using a numeric keypad (BUFFALO BSTK100BK, Nagoya, Japan).

In Experiment 1, the monocular presentation was controlled using a pair of visual occlusion goggles (PLATO Visual Occlusion Spectacles; Translucent Technologies, Toronto, Canada), which allowed precise and rapid shuttering of the visual input to each eye. The participants viewed the display through the goggles, enabling the presentation of adaptation and test stimuli to either the same or the opposite eye. In Experiments 2 and 3, the participants viewed the display binocularly without wearing goggles.

### Stimuli

The visual stimuli were designed using MATLAB with the Psychtoolbox extensions (Brainard, 1997; Kleiner et al., 2007; Pelli, 1997). All stimuli were presented in white (104.8 cd/m^2^) against a medium-gray background (55.0 cd/m^2^). A white fixation cross (0.5°) was continuously displayed at the center of the screen, except during the perceptual reporting phase, via a button press.

Three types of adaptation stimuli were used (Figure 1A, top): (1) digital numerals 6 and 8 (Experiments 1 and 2), (2) the upper parts of digital numerals 6 and 8 (hereafter termed “digital elements,” Experiment 3), and (3) white noise patterns as control stimuli (all experiments). White noise stimuli were generated by randomly shuffling the pixels of digital numeral 6. The digital numeral and white noise stimuli subtended to approximately 174 pixels in height (approximately 5.1° in visual angle), whereas the digital element stimuli used in Experiment 3 were half this height. The width of all the stimuli was 98 pixels (approximately 2.9°).

**Figure 1.**
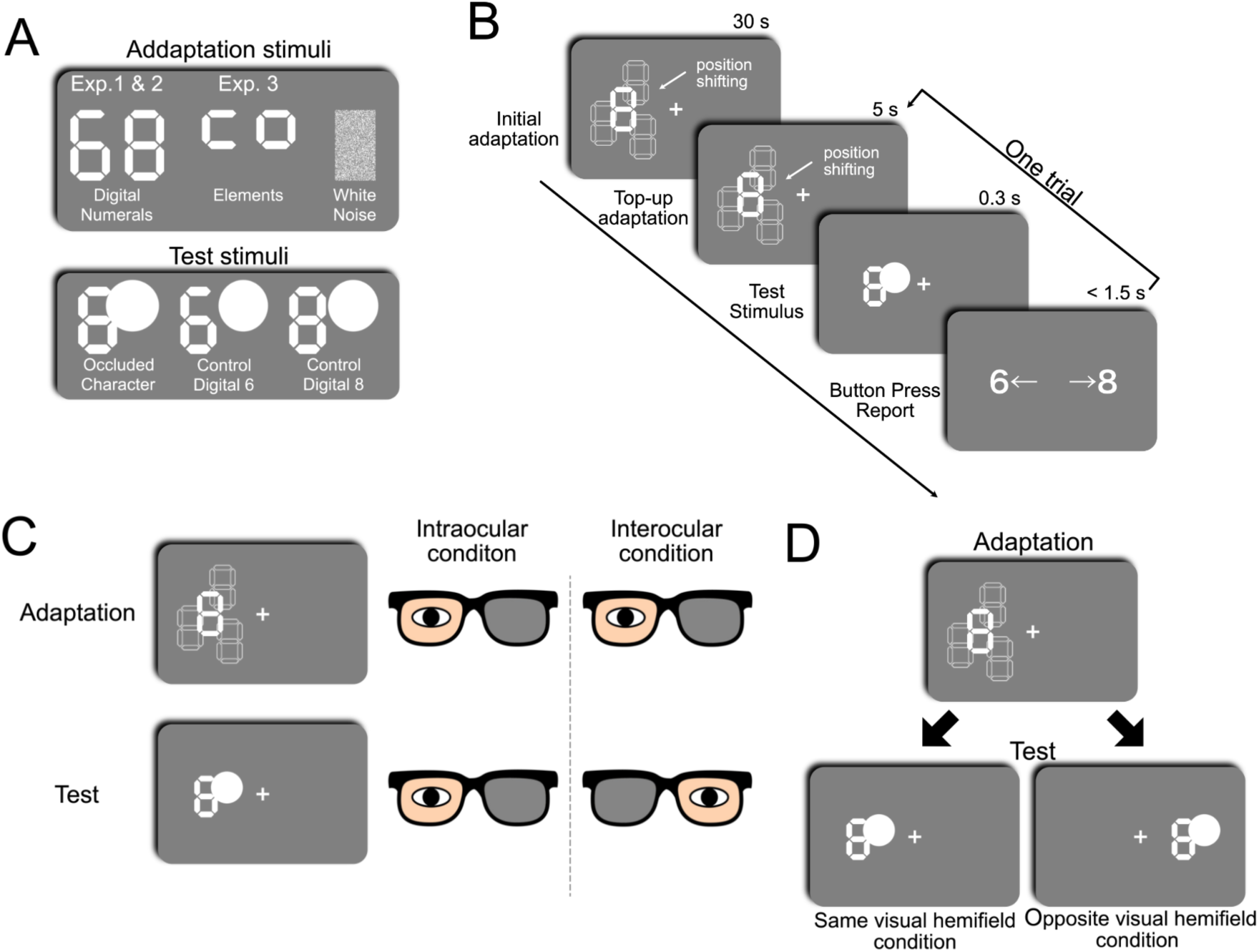
Experimental design. (**A**) Stimuli used in the experiment. *Top panel:* three types of visual stimuli were used as adaptation stimuli: digital numerals (for Experiments 1 and 2), digital elements (shape corresponding to the upper part of digital numerals, for Experiment 3), and control stimuli (white noise, all experiments). *Bottom panel:* three types of test stimuli were used: an occluded numeral (with a circular mask covering the upper-right portion) and two control numerals presented with a nonoverlapping mask. (**B**) Time course of the experimental procedure. Each block began with an initial adaptation phase in which adaptation stimuli were presented for 30 s, with their position shifted every 1 s to minimize low-level adaptation. This was followed by a sequence of trials. Each trial consisted of a 5-s top-up adaptation using the same adaptation stimuli, a 300-ms test stimulus presentation, and a response period. The participants indicated whether they perceived the test stimulus as 6 or 8 by pressing the corresponding key on a numeric keypad. (**C**) Schematic illustrations of the ocular conditions in Experiment 1 using the visual occlusion goggle device. In the intraocular condition (*middle*), adaptation (*top*) and test stimuli (*bottom*) were presented to the same eye. In the interocular condition (*right*), adaptation and test stimuli were presented to opposite eyes. (**D**) Visual hemifield conditions were tested in Experiments 2 and 3. In the same visual hemifield condition (*left*), adaptation (*top*) and test stimuli (*bottom*) were presented in the same visual hemifield. In the opposite visual hemifield condition (*right*), adaptation and test stimuli were presented in opposite visual hemifields across the vertical meridian.

We used three types of test stimuli (Figure 1A, bottom). The main test stimulus was a digital numeral partially occluded by a white circular mask placed over its upper-right portion (Luo et al., 2024). Two control test stimuli consisting of unoccluded digital numerals (6 or 8) were presented with a nonoverlapping circular mask positioned in the upper-right area. All test stimuli matched the size of the adaptation digital numerals. The circular mask had a diameter of 153 pixels (approximately 4.5°). The control test stimuli were included to confirm task engagement and ensure the reliability of the perceptual responses to partially occluded test stimuli. For this control test stimulus, the participants responded correctly in 96.92% (standard deviation [SD] across participants = 0.0271), 98.2% (SD = 0.0151), and 98.92% (SD = 0.0075) of the trials in Experiments 1, 2, and 3, respectively, confirming that their perceptual reports of the partially occluded stimuli were reliable.

Figure 1B illustrates the temporal sequence of the trial, similar to that used by Luo et al. (2024). Adaptation and test stimuli were presented in either the left or right visual hemifield. The adaptation stimuli were presented at an eccentricity of 6° from the fixation point. The positions were randomly shifted once per second within a circular region (diameter, 6°). This positional shifting was intended to minimize low-level adaptation in early visual areas (e.g., V1), where receptive fields are relatively small, and ensure that the stimuli remained outside the fovea. Test stimuli were presented at 6° eccentricity and along the horizontal meridian but remained static throughout the presentation.

### Procedure and experimental conditions

Participants were instructed to maintain fixation on the central white cross throughout the session. Before the trial sequence began, an initial adaptation phase of 30 s was performed (Figure 1B; see Supplementary Movies S1–S8). This was followed by a 0.5-s blank period during which only the fixation cross was presented. Each subsequent trial consisted of a 5-s top-up adaptation to the same stimulus, a 0.1-s blank period, and a 0.3-s test stimulus presentation. Upon the disappearance of the test stimulus, participants reported whether they perceived it as 6 or 8 by pressing the corresponding keys on a numeric keypad. A 0.5-s inter-trial interval with the fixation cross preceded the top-up adaptation phase of the next trial.

Each experiment consisted of 12 blocks: four blocks with adaptation stimulus to the numeral 6 (or its digital element), four blocks with adaptation to the numeral 8 (or its digital element), and four blocks with the white noise pattern used as a control condition. Within each block, the three test stimuli (Figure 1A, right) were presented 20 times in random order, yielding 60 trials per block.

This design resulted in 80 repetitions for each adaptation-test stimulus pair across the experiment, for a total of 720 trials (60 trials × 12 blocks). Each block lasted approximately 7–8 min, and the participants were allowed to take short breaks between blocks. The full experimental session lasted approximately 120 min.

### Experimental conditions in Experiment 1

In Experiment 1, the participants were adapted to digital numerals (6 or 8) or white noise (control). The adaptation and test stimuli were always presented in the same visual hemifield. The visual hemifield for the stimulus presentation (left or right) was randomly selected for each block. Two ocular conditions were tested (Figure 1C). In the intraocular condition, the adaptation and test stimuli were presented to the same eye, whereas in the interocular condition, the stimuli were presented to different eyes. Across the 12 blocks, six were assigned to the intraocular condition and six to the interocular condition. Block order was randomized without counterbalancing the two ocular conditions. However, the adapted eye alternated across blocks to ensure equal stimulation of the left and right eyes.

### Experimental conditions in Experiment 2

In Experiment 2, the participants were adapted to digital numerals (6 or 8) or white noise (control). Each adaptation condition consisted of four blocks divided equally between the left and right visual hemifields (two blocks per hemifield). Across the 12 blocks, the hemifield of the stimulus presentation alternated between blocks, with the starting hemifield counterbalanced across the participants. Two visual hemifield conditions were tested (Figure 1D). In the same visual hemifield condition, the adaptation and test stimuli appeared in the same visual hemifield, whereas in the opposite visual hemifield condition, the two stimuli were presented in opposite hemifields. Within each block (60 trials), half of the trials corresponded to each condition in randomized order.

### Experimental conditions in Experiment 3

In Experiment 3, the participants were adapted to the digital elements (6 or 8) or white noise (control). The same two hemifield conditions as in Experiment 2 (same vs. opposite visual hemifield) were tested. Other procedures were the same as those used in Experiment 2.

### Statistical analyses and quantification

To evaluate the effect of adaptation on the bistable perception of partially occluded digital numerals, we calculated the proportion of trials in which the participants reported the occluded digital numeral as either 6 or 8 for each adaptation condition. The perceptual bias was quantified using the following bias index:

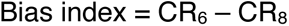

where *CR_6_* and *CR_8_* denote the choice rates for 6 or 8 responses, respectively. The index ranged from −1 to 1, with positive values indicating a bias toward perceiving 6 and negative values indicating a bias toward perceiving 8.

The distribution of bias indices across the participants using violin plots is shown in Figures 2–4 (Bechtold et al., 2021; https://zenodo.org/records/4559847). Differences in bias indices between the adaptation conditions were assessed using the Wilcoxon signed-rank test. The effect sizes were quantified using a rank-biserial correlation (presented as r in the Results). In addition, robust paired Cohen’s d was reported as a metric of effect size for reference.

**Figure 2.**
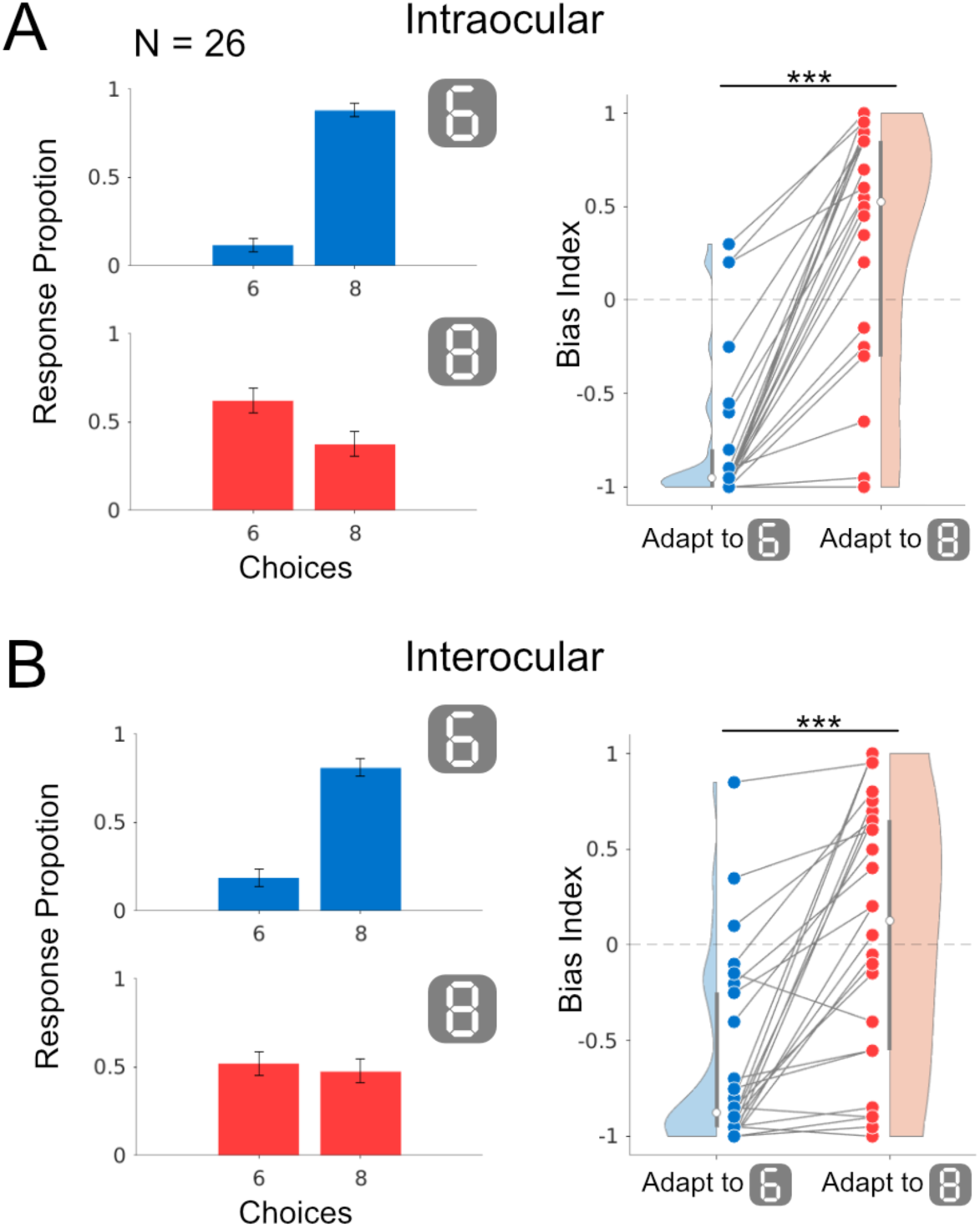
Results of Experiment 1. (**A**) Intraocular condition. *Left panel*: Response proportions of the participants reporting 6 or 8 after adaptation to digital numerals 6 (*top*, blue) and 8 (*bottom*, red). Strong, yet opposite, perceptual biases were observed across two adaptation conditions. Error bars indicate ± 1 standard error (SE) from the mean. *Right panel*: Bias indices from 26 participants. Consistent with the response proportions, the distributions of bias indices shifted in opposite directions for the two adaptation stimuli (blue, adaptation to digital numeral 6; red, adaptation to digital numeral 8). Individual data points are shown as dots, with paired data connected by gray lines. Shaded regions represent kernel density estimates. White circles indicate the median across participants, and the black vertical lines denote ± 1 standard deviation (SD) from the mean. (**B**) Interocular condition. The perceptual bias, similar to the intraocular condition, was observed, although with a smaller effect size. Conventions are identical to those used in panel A. Asterisks indicate statistical significance (Wilcoxon signed-rank test, ***p < 0.001).

**Figure 3.**
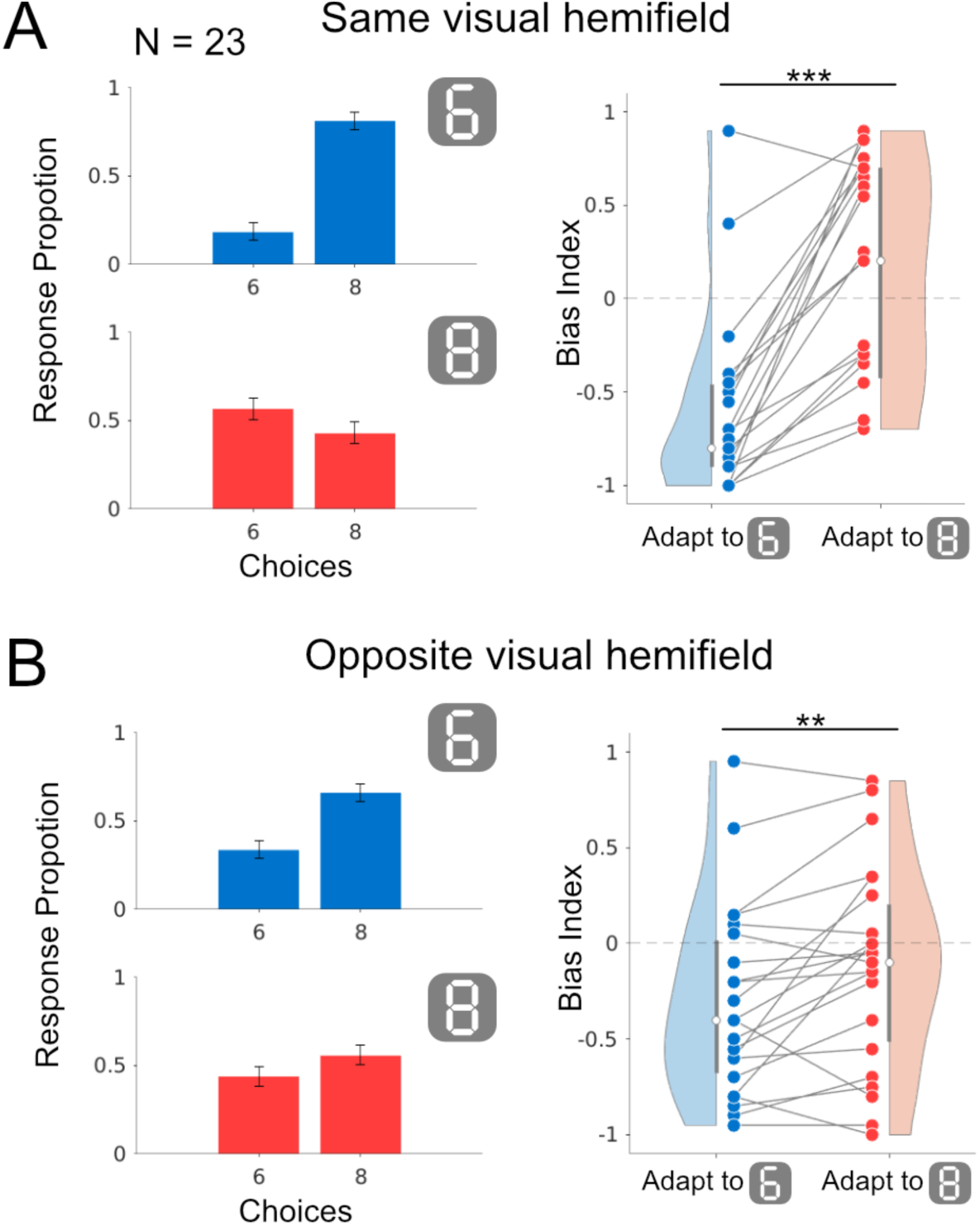
Results of Experiment 2. (**A**) Same visual hemifield condition. *Left panel*: Response proportions of the participants reporting 6 or 8 after adaptation to digital numerals 6 (*top*, blue) and 8 (*bottom*, red). *Right panel*: Bias indices from 23 participants. Bias indices significantly shifted in opposite directions for the two adaptation stimuli (blue, adaptation to digital numeral 6; red, adaptation to digital numeral 8). Other conventions are identical to those used in Figure 2. (**B**) Opposite visual hemifield condition. Significant perceptual bias induced by adaptation was observed, although of a smaller effect size than in the same visual field condition. Asterisks indicate statistical significance (Wilcoxon signed-rank test, **p < 0.01, ***p < 0.001).

**Figure 4.**
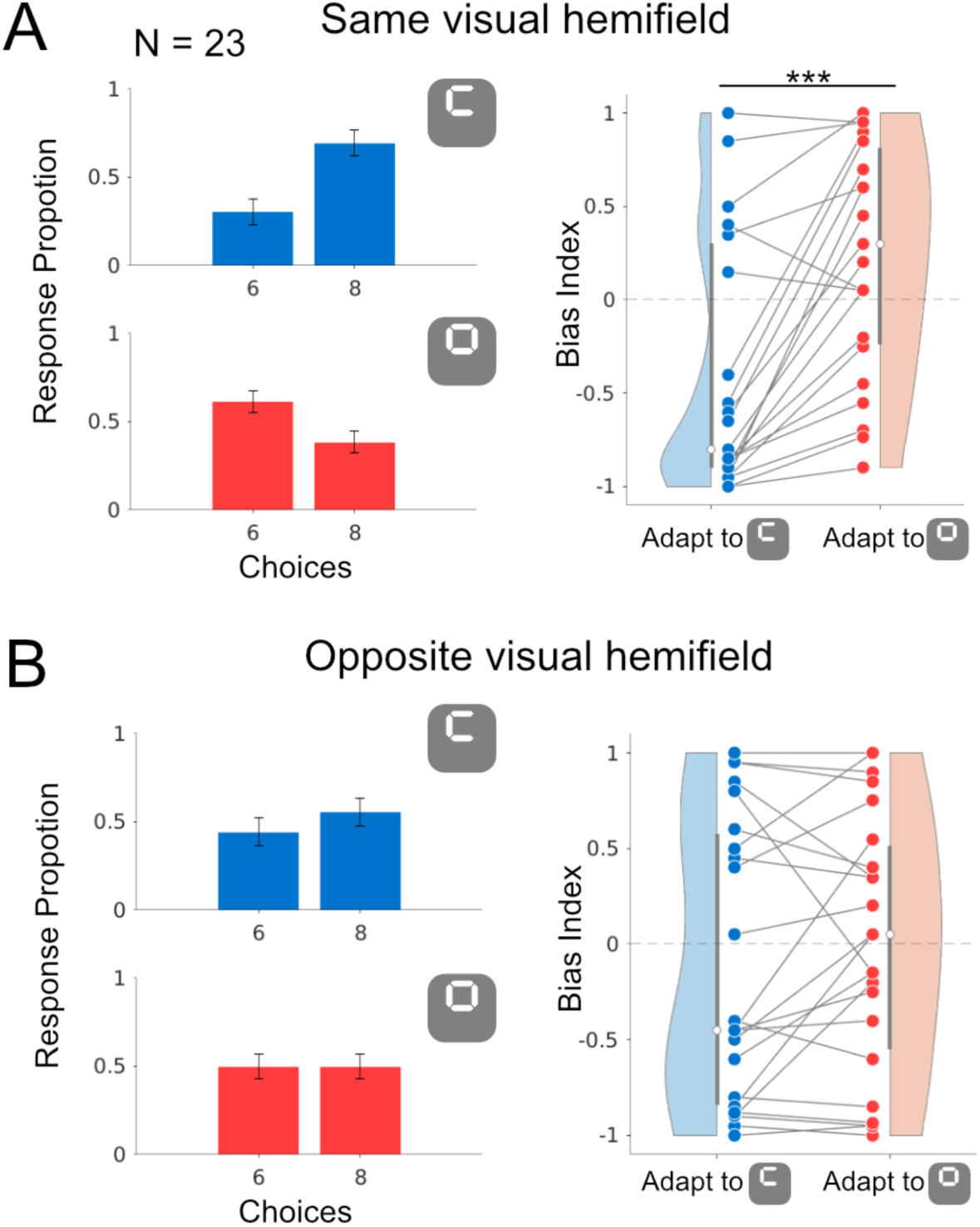
Results of Experiment 3. (**A**) Same visual hemifield condition. *Left panel*: Response proportions of the participants reporting 6 or 8 after adaptation to the digital element of 6 (*top*, blue) and 8 (*bottom*, red). *Right panel*: Bias indices from 23 participants (blue, adaptation to the upper element of digital numeral 6; red, adaptation to the upper element of digital numeral 8). Other conventions are identical to those used in Figure 2. Although weaker than in the digital numeral conditions (Figure 3), perceptual biases in the opposite direction were still observed across the two adaptation conditions. (**B**) Opposite visual hemifield condition. In this condition, the differences in bias indices between the two adaptation conditions did not reach statistical significance. Asterisks indicate statistical significance (Wilcoxon signed-rank test, ***p < 0.001).

### Code availability

Data and code for generating visual stimuli and replicating figures in this work are available on GitHub upon acceptance of the manuscript (https://github.com/OkazakiTakemuraLab/SymbolicNumeralAdaptationTransfer).

## Results

Here, we report the results of three experiments examining the extent to which the symbolic numeral adaptation effect is transferred across the eyes and visual hemifields.

### Experiment 1

Figure 2 shows the results of Experiment 1, which tested symbolic numeral adaptation under two conditions: intraocular (adaptation and test stimuli were presented to the same eye) and interocular (adaptation and test stimuli were presented to the opposite eye).

Under intraocular conditions (Figure 2A), adaptation induced a significant perceptual bias, which is consistent with a previous study (Luo et al., 2024). After adapting to numeral 6, the participants more often reported the ambiguous test stimulus as 8 (Figure 2A, *left top*; mean response proportions: 11.7% for 6; 88.3% for 8), whereas adaptation to 8 led to more reports of 6 (Figure 2A, *left bottom*; 62.4% for 6, 37.6% for 8). Bias indices (see Methods) confirmed these effects: adaptation to 6 produced a mean bias index of –0.77, significantly different from that produced by adaptation to 8 (mean = 0.248, r = –1.00, d = –1.92; Wilcoxon signed-rank test, p = 0.00003), demonstrating strong adaptation within the same eye.

Similar patterns were observed under the interocular condition (Figure 2B). Following adaptation to 6, the participants reported 8 more frequently (Figure 2B, *left top*; 18.8% for 6; 81.3% for 8), and following adaptation to 8, 6 was reported more often (Figure 2B, *left bottom*; 52.2% for 6, 47.8% for 8). The corresponding bias indices averaged across the participants were –0.63 and 0.04, respectively, and the difference was statistically significant (r = –0.92, d = –0.96, p = 0.00004), although the effect size was smaller than that of the intraocular condition. Notably, in both the intraocular and interocular conditions, adaptation to white noise did not induce a significant perceptual bias (Supplementary Figure 1).

These results indicate that symbolic numeral adaptation can be transferred across eyes, suggesting that the underlying mechanisms involve visual processing stages after binocular integration.

### Experiment 2

Figure 3 shows the results of Experiment 2, which assessed the transfer of symbolic numeral adaptation across visual hemifields. Two hemifield conditions were tested: the same visual hemifield condition (adaptation and test stimuli were presented in the same visual hemifield) and the opposite visual hemifield condition (adaptation and test stimuli were presented in the opposite visual hemifield across the vertical meridian [VM]).

Under the same visual hemifield condition (Figure 3A), adaptation produced significant perceptual biases, consistent with those reported in a previous study (Luo et al., 2024). After adapting to numeral 6, the participants more often reported 8 (Figure 3A, *left top*; 18.6% for 6; 81.4% for 8), whereas after adapting to 8, they more frequently reported 6 (Figure 3A, *left bottom*; 56.9% for 6, 43.2% for 8). Bias indices confirmed this effect: adaptation to 6 yielded a mean index of –0.63, which was significantly different from the adaptation to 8 (mean = 0.14, r = –0.99, d = -1.39, p = 0.00003), demonstrating a strong adaptation within the same visual hemifield.

In the opposite visual hemifield condition (Figure 3B), similar biases were observed but with a reduced magnitude. After adapting to 6, the participants more often reported 8 (Figure 3B, *left top*; 33.9% for 6; 66.1% for 8). In contrast, adaptation to 8 did not lead to more reports of 6, although the proportion of reports of 8 was lower than that observed after adaptation to 6 (Figure 3B, *left bottom*; 44.0% for 6, 56.0% for 8). The bias indices were –0.32 and –0.12, respectively; this difference was statistically significant, although the effect size was modest, and notable individual differences were present (r = –0.65, d = –0.42, p = 0.007). Similar to Experiment 1, these systematic perceptual biases were not observed when white noise was used as the control stimulus (Supplementary Figure 2) in either the same or opposite visual hemifield conditions.

These results indicate that symbolic numeral adaptation can be transferred across visual hemifields, although the effect is attenuated when adaptation and test stimuli are presented in opposite hemifields.

### Experiment 3

Figure 4 shows the results of Experiment 3, which tested whether adaptation to digital elements (upper parts of digital numerals) was transferred across the hemifields.

In the same visual hemifield condition (Figure 4A), adaptation to the digital element of 6 led participants to report 8 more often (Figure 4A, *left top*; 30.4% for 6 vs. 69.6% for 8), whereas adaptation to the digital element of 8 led to more reports of 6 (Figure 4A, *left bottom*; 61.5% for 6 vs. 38.6% for 8). Bias indices confirmed the significant difference: adaptation to 6 produced a mean index of –0.39, compared with 0.23 after adaptation to 8 (r = –0.89, d = –0.83, p = 0.0002). This effect was consistent with that reported in a previous study (Luo et al., 2024).

In the opposite visual hemifield condition (Figure 4B), no significant adaptation effects were observed. After adaptation to the element of 6, the participants more often reported 8 (Figure 4B, *left top*; 44.4% for 6 vs. 55.6% for 8), whereas after adaptation to the element of 8, responses were evenly distributed (Figure 4B, *left bottom*; 50.0% for 6 vs. 50.0% for 8). The bias indices were –0.11 and 0.0007, respectively, and the difference was not statistically significant (r = –0.17, d = –0.13, p = 0.38). As in the other two experiments, adaptation to white noise did not induce a systematic perceptual bias in either hemifield condition (Supplementary Figure 3).

These results suggest that adaptation to the digital elements can bias perception within the same visual hemifield, but provide no evidence of transfer across the VM.

## Discussion

In this study, we conducted three psychophysical experiments to test whether adaptation effects on symbolic numerals and their shape components could be transferred between the eyes or across visual hemifields. In Experiment 1, adaptation to digital numerals induced perceptual biases even when the adaptation and test stimuli were presented to different eyes, indicating an interocular transfer of the adaptation effect. In Experiment 2, adaptation to digital numerals also induced a small but statistically significant perceptual bias when the adaptation and test stimuli were presented in the opposite visual hemifields, supporting the existence of a transfer of the adaptation effect across hemifields. Finally, in Experiment 3, when the adaptation stimuli consisted of parts of digital numerals, no significant perceptual bias was observed when the adaptation and test stimuli were presented in the opposite visual hemifields. Below, we discuss the implications of these findings and their relationship to the literature.

### Symbolic numeral adaptation exhibited interocular transfer

In general, adaptation effects involving relatively higher-level visual processing transfer between eyes (Banks et al., 1975; Kompaniez et al., 2013; Mitchell & Ware, 1974; Movshon et al., 1972; Nishida et al., 1994; Raymond, 1993; Steiner et al., 1994). Physiologically, a traditional model holds that visual information from the two eyes is processed separately at an earlier stage of visual processing (Hubel & Wiesel, 1972), although the physiological locus of binocular integration remains an active topic of investigation (Maier et al., 2022). This finding supports the prevailing view that monocular processing occurs earlier than binocular integration.

The results from Experiment 1 (Figure 2) revealed a significant interocular transfer of the adaptation effect for symbolic numerals, indicating that the effect arises at or beyond the stages of visual processing where binocular integration has occurred (Figure 5).

**Figure 5.**
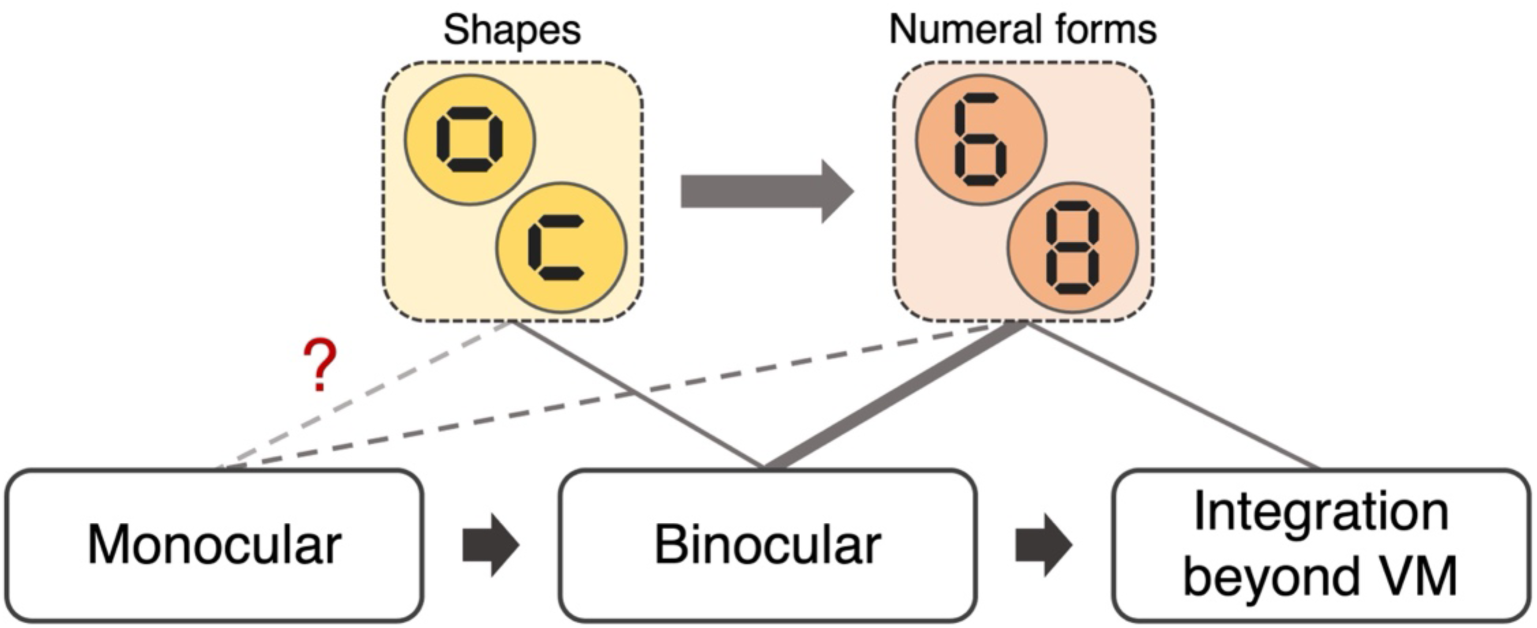
Schematic model of hierarchical visual information processing and results of this study. The visual information processing is composed of early monocular, binocular, and later processing, including integration of visual information beyond the vertical meridian (VM). Our results suggest that perceptual bias induced by adaptation to the digital elements (shapes) and digital numerals (numeral forms) is associated with each processing stage differently. The schematic strength of association is depicted by the thickness of lines. Notably, the involvement of the monocular processing stage for digital elements remains uncertain as we did not perform a comparison between intraocular and interocular transfer for these adaptation stimuli.

Notably, the magnitude of the adaptation effect in the interocular condition was smaller than that in the intraocular condition, although the interocular transfer remained significant (Figure 2). This difference suggests the partial contribution of monocular processing to adaptation. Alternatively, other factors may account for the differences in magnitude between interocular and intraocular conditions. Specifically, the interocular condition might have hindered task performance because prolonged exposure to a blank screen without dynamic input to one eye could reduce the responsiveness of the visual system or because the brief presentation of the test stimulus in the other eye might make the task more attentionally demanding. Therefore, these results may not provide definitive evidence for the strict involvement of monocular processing in the adaptation effect, although these multiple possible explanations are not mutually exclusive.

### Symbolic numeral adaptation transfers to the opposite visual hemifield

Psychophysically, some higher-order adaptation effects transfer across visual hemifields (Meng et al., 2006; Nakayama et al., 2024; Palmer & Clifford, 2022). Physiologically, although the early visual areas primarily receive input from the contralateral visual hemifield, the intraparietal sulcus and higher visual field maps increasingly represent the ipsilateral field (Amano et al., 2009; Mackey et al., 2017; Tootell et al., 1998). This gradual increase in the ipsilateral representation is likely mediated by callosal connections that transmit information across the VM. Crucially, such connections are known to be absent in the horizontal visual hemifield representation of V1 (Caspers et al., 2015; Dougherty et al., 2005; Van Essen & Zeki, 1978). Therefore, the integration of information across the VM, specifically regarding stimuli presented around the horizontal meridian, is likely to occur at the intermediate to higher stages of visual processing.

The present finding that the adaptation effect transferred across visual hemifields for test stimuli presented at the horizontal meridian suggests the involvement of higher-level processing stages, where interhemispheric integration of visual information takes place (Figure 5). However, the smaller effect size observed in the opposite visual hemifield condition compared with that in the same visual hemifield condition indicates that such a higher-level processing stage alone cannot fully explain the adaptation effect. Instead, it is likely that the lower or intermediate processing stages before interhemispheric integration also contribute to the observed adaptation effects.

### Adaptation occurring at the shape processing stage

A previous study (Luo et al., 2024) demonstrated that adaptation effects were induced even when participants were exposed only to the elemental parts of digital numerals. This finding suggests that mid-level shape processing mechanisms, which precede number form processing, may partly contribute to the symbolic numeral adaptation phenomenon. However, adaptation at this stage alone cannot fully account for the overall effect of symbolic numeral adaptation, as the effect size becomes smaller than that observed with adaptation to complete symbolic numerals (Luo et al., 2024).

We replicated this finding under the same visual hemifield conditions, confirming that adaptation to digital elements can bias the perception of occluded numerals (Experiment 3, Figure 4). However, we found no statistically significant transfer across visual hemifields. Therefore, shape-level adaptation may primarily occur at a relatively early stage of visual processing, specifically prior to the stage at which information from the left and right visual hemifields is integrated (Figure 5).

### Open questions and future directions

In Experiments 1 and 2, we observed that adaptation to 6 tended to yield a more robust perceptual bias toward 8, whereas adaptation to 8 resulted in a relatively weaker bias (Figures 2 and 3). This asymmetry was more pronounced in the opposite visual hemifield condition of Experiment 2, where the overall effect size was smaller and adaptation to 8 did not induce a clear bias toward 6. Although speculative, these findings suggest that the impact of symbolic numeral adaptation may be asymmetric across numerals (e.g., 6 vs. 8). The mechanism underlying this nonidentical adaptation effect remains unclear.

Specifically, it is worth considering whether such an asymmetry reflects differences in lifetime exposure frequency (Benford, 1938), although addressing this possibility is beyond the scope of this study.

In conjunction with those of a previous study (Luo et al., 2024), our findings suggest that the symbolic numeral adaptation effect involves middle- to higher-level stages of visual processing. Although we cannot directly identify the neural substrates of this phenomenon solely based on the results of this study, several plausible cortical areas may be considered. One possibility is the visual number form area, which exhibits selective responses to numerical symbols (Abboud et al., 2015; Grotheer et al., 2016; Shum et al., 2013). Another candidate is a region in the temporal-occipital cortex (NTO; Cai et al., 2023), which has also shown symbolic numeral selectivity. In addition, some neuroimaging studies have reported involvement of regions in the intraparietal sulcus in symbolic numeral processing, using functional magnetic resonance imaging adaptation or meta-analysis (Piazza et al., 2007; Sokolowski et al., 2017; Vogel et al., 2015). A direct investigation of how these areas are involved in the symbolic numeral adaptation effect requires neuroimaging or electrophysiological measurements. Future studies may provide further insights into how the subjective recognition of symbolic numerals emerges during visual processing in the brain.

## Conclusion

The symbolic numeral adaptation effect is transferred across both eyes and hemifields. In contrast, we did not find evidence of interhemifield transfer for the adaptation effect induced by element stimuli targeting the shape processing stage. Taken together, these findings suggest that the subjective perception of symbolic numerals depends on the middle- to higher-level visual processing stages, where information is integrated across the eyes and hemifields.

## Acknowledgements

We thank Kumiko Kobayashi and Yuri Morii for their support in data curation and participant recruitment. This study was supported by the Japan Society for the Promotion of Science (JSPS) KAKENHI (JP24K03240 to H.T.). We thank Editage (http://www.editage.com) for editing and reviewing this manuscript for the English language.

## Supplementary Figures

**Supplementary Figure 1.**
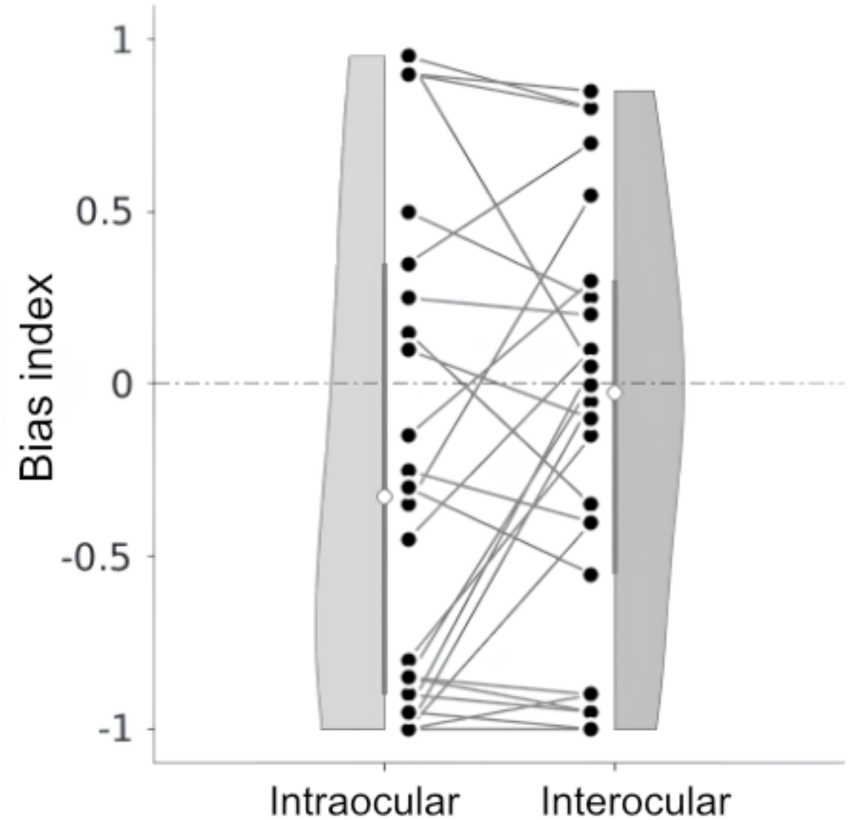
Results of the control condition in Experiment 1. Bias indices from 26 participants under the control adaptation conditions, in which the participants adapted to white noise rather than digital numerals. The distribution of bias indices (vertical axis) was comparable between the two ocular conditions (left, intraocular; right, interocular). Dots represent individual participants, with paired data connected by gray lines. White circles indicate the median across participants. Shaded areas indicate kernel density estimates of the distribution. Error bars indicate ± 1 SD from the mean.

**Supplementary Figure 2.**
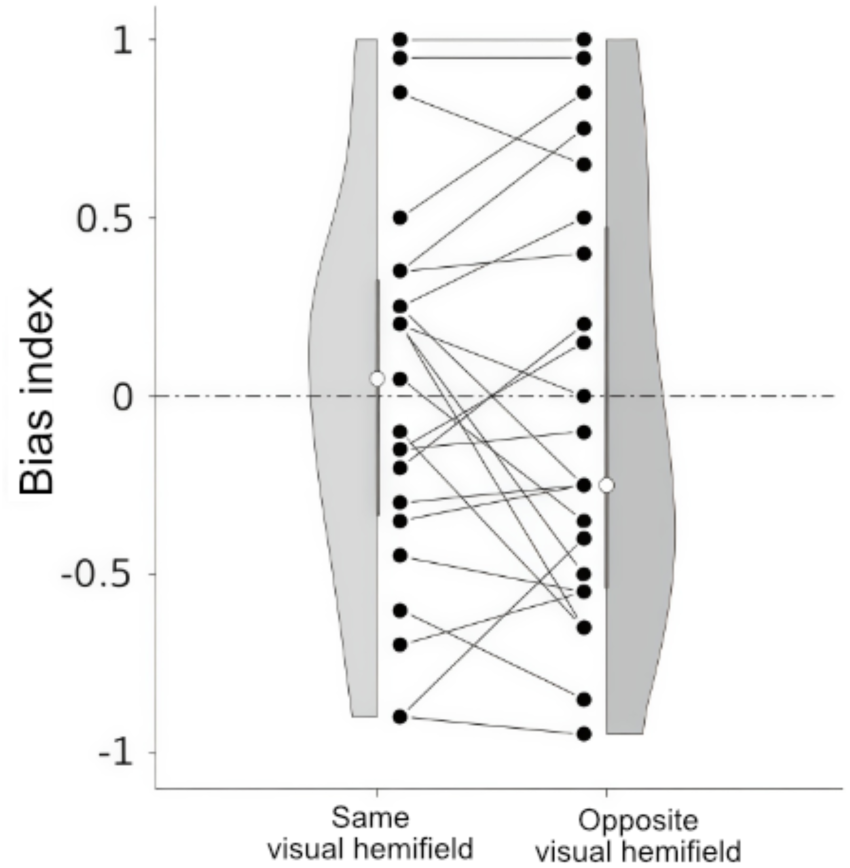
Results of the control condition in Experiment 2. Bias indices from 23 participants under the control adaptation conditions, in which the participants adapted to white noise rather than digital numerals. The distribution of bias indices (vertical axis) was comparable between the two visual hemifield conditions (left, same visual hemifield; right, opposite visual hemifield). Conventions are identical to those in Supplementary Figure 1.

**Supplementary Figure 3.**
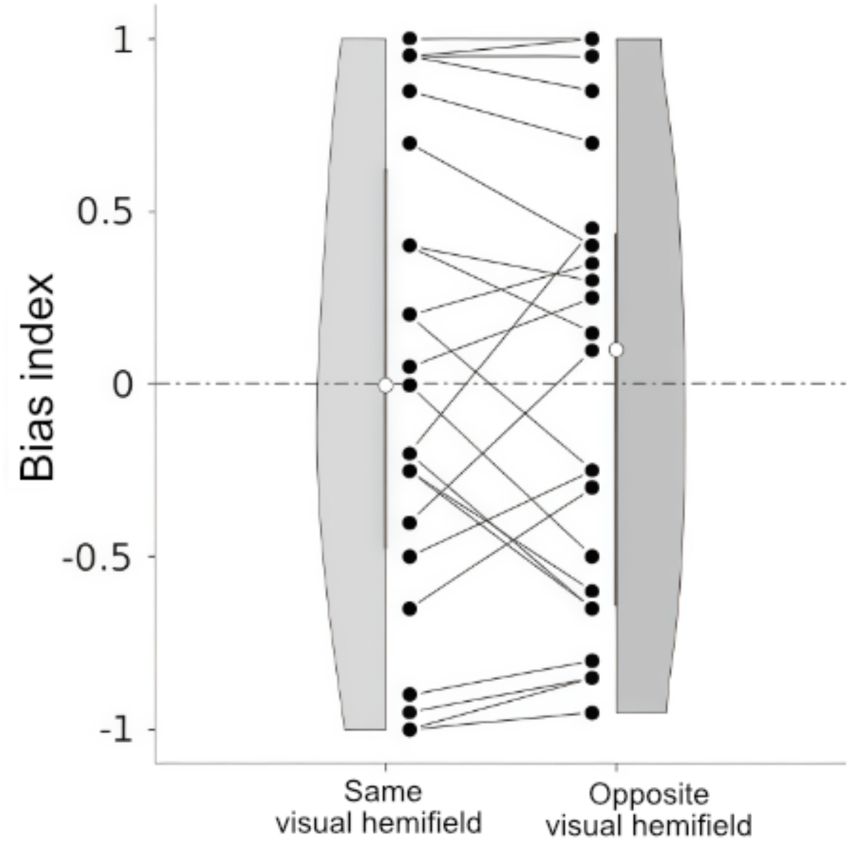
Results of the control condition in Experiment 3. Bias indices from 23 participants under the control adaptation conditions, in which participants adapted to white noise rather than digital elements. The distribution of bias indices was comparable between the two visual hemifield conditions (left, same visual hemifield; right, opposite visual hemifield). Conventions are identical to those in Supplementary Figure 1.

**Supplementary Movie 1.**

Demonstration video of the stimulus presentation sequence corresponding to the same visual hemifield conditions in Experiments 2. The adaptation stimulus (the digital numeral 6, presented in the left visual hemifield) is shown for 30 s, followed by the occluded test stimulus presented for 0.3 s in the same visual hemifield.

**Supplementary Movie 2.**

Demonstration video of the stimulus presentation sequence corresponding to the opposite visual hemifield in Experiment 2. The adaptation stimulus (the digital numeral 6, presented in the left visual hemifield) is shown for 30 s, followed by the occluded test stimulus presented for 0.3 s in the opposite (right) visual hemifield.

**Supplementary Movie 3.**

Demonstration video of the stimulus presentation sequence corresponding to the same visual hemifield conditions in Experiments 2. The adaptation stimulus (the digital numeral 8, presented in the left visual hemifield) is shown for 30 s, followed by the occluded test stimulus presented for 0.3 s in the same visual hemifield.

**Supplementary Movie 4.**

Demonstration video of the stimulus presentation sequence corresponding to the opposite visual hemifield in Experiment 2. The adaptation stimulus (the digital numeral 8, presented in the left visual hemifield) is shown for 30 s, followed by the occluded test stimulus presented for 0.3 s in the opposite visual hemifield.

**Supplementary Movie 5.**

Demonstration video of the stimulus presentation sequence corresponding to the same visual hemifield conditions in Experiments 3. The adaptation stimulus (the digital element of 6, presented in the left visual hemifield) is shown for 30 s, followed by the occluded test stimulus presented for 0.3 s in the same visual hemifield.

**Supplementary Movie 6.**

Demonstration video of the stimulus presentation sequence corresponding to the opposite visual hemifield in Experiment 3. The adaptation stimulus (the digital element of 6, presented in the left visual hemifield) is shown for 30 s, followed by the occluded test stimulus presented for 0.3 s in the opposite (right) visual hemifield.

**Supplementary Movie 7.**

Demonstration video of the stimulus presentation sequence corresponding to the same visual hemifield conditions in Experiments 3. The adaptation stimulus (the digital element of 8, presented in the left visual hemifield) is shown for 30 s, followed by the occluded test stimulus presented for 0.3 s in the same visual hemifield.

**Supplementary Movie 8.**

Demonstration video of the stimulus presentation sequence corresponding to the opposite visual hemifield in Experiment 3. The adaptation stimulus (the digital element of 8, presented in the left visual hemifield) is shown for 30 s, followed by the occluded test stimulus presented for 0.3 s in the opposite visual hemifield.

